# An Integrated EEG, TMS-EEG and Behavioural Dataset for Investigating the Neural Correlates of Working Memory

**DOI:** 10.64898/2026.07.12.738085

**Authors:** Mana Biabani, Alex Fornito, Sarah Thompson, Ziarih Hawi, Murat Yucel, Mark A. Bellgrove, Nigel C. Rogasch

## Abstract

Working memory is a fundamental process that underlies cognition. Accordingly, the neural mechanisms that support working memory performance are of great interest in cognitive neuroscience. We present a publicly accessible dataset comprising 123 healthy adults (mean age = 28.58 years, SD = 7.56; age range = 18–46 years; 75 females) to facilitate investigation of the neural mechanisms underlying working memory performance in humans. Across two days of assessment, participants underwent a battery of cognitive tasks assessing working memory and other cognitive domains (day 1), followed by 62-channel EEG recordings (day 2). EEG data were acquired at rest, during a visual working memory task, and following single-pulse transcranial magnetic stimulation (TMS-EEG) targeting key brain regions involved in working memory. This article provides a detailed overview of the study design, methodology, and data characteristics.

## Background and summary

Working memory is a fundamental cognitive function that supports a broad range of higher order processes such as reasoning, emotional regulation, and social behaviour (Baddeley, 2012; Cowan, 2014). Visual working memory (VWM), the ability to hold and manipulate visual information over short periods, plays a critical role in cognitive function. Differences in VWM capacity account for over 40% of the variance in cognitive ability across both healthy individuals and those with cognitive impairments (e.g., schizophrenia and advanced age) (Luck and Vogel, 2013). Remarkably, this cognitive construct can be accurately assessed using simple tasks like recalling coloured squares. Evidence shows that memory for such basic stimuli is highly predictive of broader cognitive abilities and is often disrupted in various neuropathological conditions (D’Esposito and Postle, 2015; Kahana and Wagner, 2024). The simplicity of VWM tasks makes them ideal for neuroimaging experiments and computational modeling, offering valuable insights into complex cognitive processes, and advancing our understanding of brain function. Human brain imaging studies using methods such as structural and functional MRI have consistently highlighted frontoparietal brain regions that are associated with VWM performance (Jonides et al., 2008; D’Esposito and Postle, 2015). However, the precise neurophysiological mechanisms and dynamic neural interactions that govern cognitive performance across these regions remain underexplored.

Electroencephalography (EEG) non-invasively captures neural activity from across the scalp with high temporal precision. Studies using EEG have revealed key neural correlates of VWM capacity including: modulation of theta (4-7 Hz), alpha (8-12 Hz), beta (15-30 Hz) and gamma (>30 Hz) band oscillations (Roux and Uhlhaas, 2014; Miller et al., 2018; Pavlov and Kotchoubey, 2022); long-range synchronisation of oscillations between regions (Fell and Axmacher, 2011); nested coupling of oscillations such as theta-gamma coupling (Lisman and Jensen, 2013); modulation of non-oscillatory activity (Voytek et al., 2015; Donoghue et al., 2020); and sustained, slow-wave event-related potentials (ERPs) such as contralateral delay activity (Fukuda et al., 2015; Luria et al., 2016; Bae and Luck, 2018). However, several key questions remain. First, it is unclear whether the EEG mechanisms recorded during a specific task generalise to broader measures of working memory performance and other cognitive domains. One approach to address this question is to model the latent structure of working memory performance using a battery of behavioural tasks (e.g., using structural equation modelling) and then correlate this latent variable with the EEG measure of interest. While this approach has proven useful for assessing the relationship between certain EEG measures and working memory performance (Unsworth et al., 2015; Lenartowicz et al., 2021), a challenge is the large populations required for this approach (usually at least 100 participants). Therefore, large datasets containing both EEG measures and broad batteries of cognitive tasks are required.

A second limitation is that the spatial resolution of EEG restricts its ability to localize the neural mechanisms underlying working memory to specific cortical regions. To address this limitation, transcranial magnetic stimulation (TMS) can be combined with EEG to non-invasively stimulate cortical regions of interest and record the resulting TMS-evoked potentials (TEPs). When applied while participants are at rest, TEPs provide unique insights into the reactivity, excitation/inhibition balance, oscillatory profile, and functional connectivity of the targeted cortical region, thereby advancing our understanding of both localized cortical function and large-scale connectivity (Ilmoniemi et al., 1997; Massimini et al., 2005; Tremblay et al., 2019). Furthermore, certain TEP features from specific cortical regions have been linked with working memory performance in healthy individuals (Rogasch et al., 2015) and those with brain disorders (Ferrarelli et al., 2012). Despite the significant advantages, TMS-EEG faces crucial challenges such as high inter-individual variability and various artifacts that can distort the signal of interest (Biabani et al., 2019, 2024; Tremblay et al., 2019; Hernandez-Pavon et al., 2023; Ziemann et al., 2026). To mitigate these issues, there is a growing call for increased data sharing of large open-access TMS-EEG datasets with rigorous experimental controls to enhance the reproducibility of TMS-EEG research (Belardinelli et al., 2019; Siebner et al., 2019).

The present dataset (Table 1) offers a unique resource including a large sample of healthy adults (n = 123) with EEG measured at rest, during a working memory task, and following single pulse TMS targeting three regions important for working memory (dorsolateral prefrontal cortex, premotor cortex, and parietal cortex) as well as a sensory control condition (shoulder stimulation). Participants also completed a battery of cognitive behavioural tests assessing working memory, response inhibition, and processing speed. We foresee many possible uses for this dataset including deep phenotyping of resting and active EEG signals to identify neural correlates of cognition, testing and optimising cleaning pipelines for EEG and TMS-EEG data, and providing a benchmark dataset for biophysical models of human neural activity at rest, during working memory and following TMS. In sum, we provide a rich, high-quality, multimodal human EEG dataset with accompanying cognitive behavioural tasks and unique concurrent TMS-EEG data. This resource will allow for reproducible investigation of neural dynamics underlying working memory with high transparency and precision.

**Table 1.** Overview of experimental design, modalities, and variables.

|  |  |
| --- | --- |
| **Design Type (s)** | • Within-subject design |
| **Measurement Type (s)** | • Brain measurements<br>• Behaviour measurements |
| **Technology Type (s)** | • Magnetic resonance imaging (MRI)<br>• Electroencephalography (EEG)<br>• Transcranial Magnetic Stimulation (TMS) |
| **Demographic and Experimental Variable (S)** | • Age<br>• Sex<br>• Education<br>• Smoker Status<br>• EEG Cap Size<br>• Distance from Screen during Task<br>• Resting Motor Threshold (rMT)<br>• Stimulation Intensity<br>• Stimulation Coordinates (x, y, z)<br>• Stimulation Condition Order<br>• Coil-to-Cortex Distance<br>• Rate of Sensory Perception during TMS<br>• Volume of White Noise during TMS<br>• Digitized Electrode Positions |
| **Sample Characteristic (s)** | • Young healthy individuals<br>• Brain |

## Methods

### Ethics approval

The research project was approved by the Monash University Human Research Ethics Committee (MUHREC), reference number 12723. The study was conducted in accordance with the Declaration of Helsinki.

### Participants

A total of 123 healthy adults (mean age = 28.58 years, SD = 7.56; age range = 18–46 years; 75 females) were tested for this study. Participants were recruited from other imaging studies run by our research group (the Genetics of Cognition [GOC] and Monash Brain Behaviour Project [MBBP]) and from the general population. Returning participants of earlier imaging studies who had consented to being contacted about future research studies were invited to participate in the current study via email. New participants were either recruited through offline advertisements at Monash University Clayton Campus, local businesses, libraries and community centres, or through online advertisements on marketing websites, social media and the university recruitment portal. Participants were initially phone screened for contraindications to TMS and MRI, colour blindness, ethnicity, and history of medical and psychiatric disorders (see Table 2). Informed consent was then obtained from participants in-person, after reading a written statement about the study and receiving a verbal explanation upon attendance. All participants consented for their de-identified data to be shared on public data repositories for use in future research. Participants were remunerated 100 AUD upon study completion.

**Table 2.**
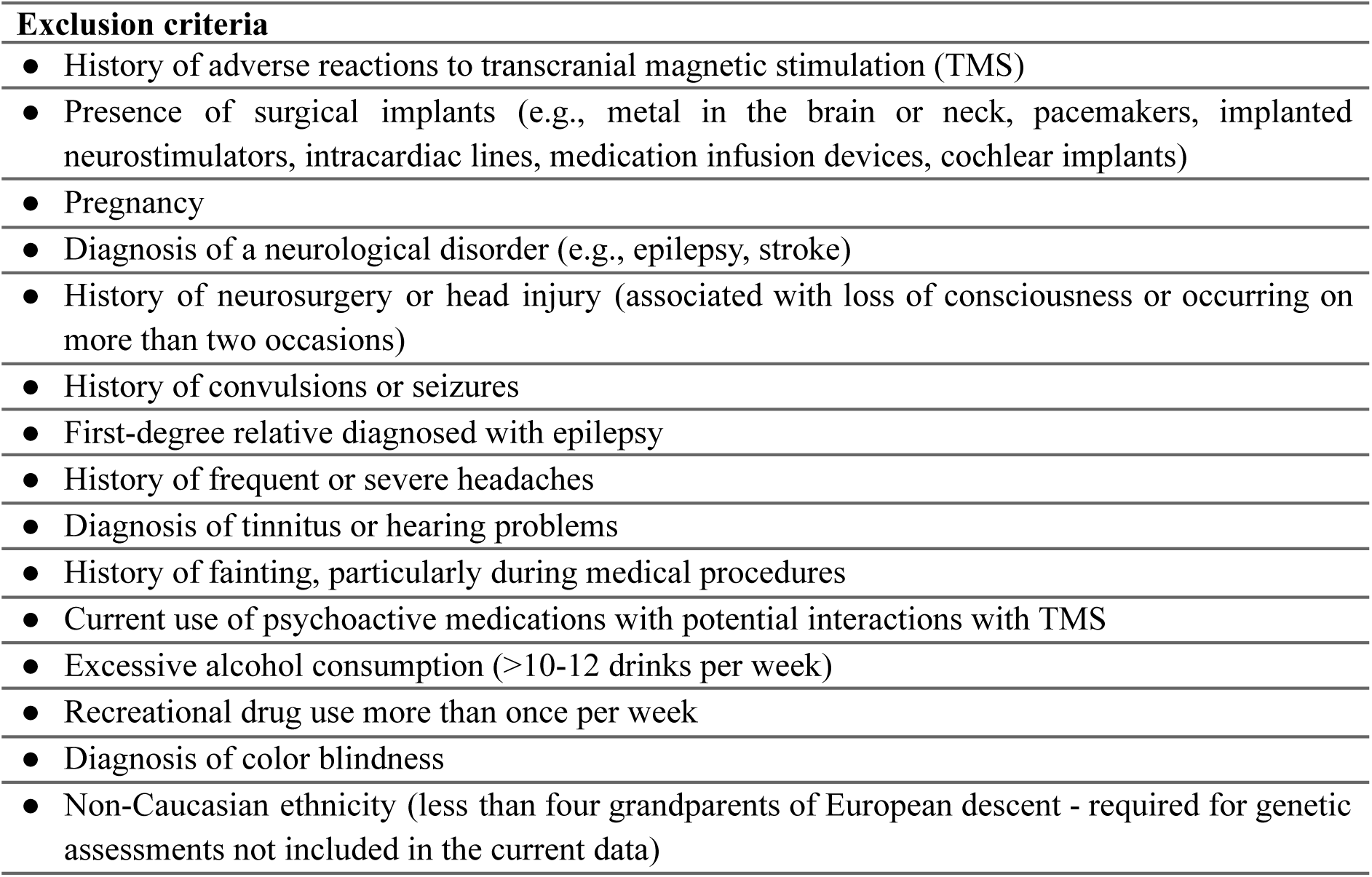
Exclusion criteria.

| **Exclusion criteria** |
| --- |
| ● History of adverse reactions to transcranial magnetic stimulation (TMS) |
| ● Presence of surgical implants (e.g., metal in the brain or neck, pacemakers, implanted neurostimulators, intracardiac lines, medication infusion devices, cochlear implants) |
| ● Pregnancy |
| ● Diagnosis of a neurological disorder (e.g., epilepsy, stroke) |
| ● History of neurosurgery or head injury (associated with loss of consciousness or occurring on more than two occasions) |
| ● History of convulsions or seizures |
| ● First-degree relative diagnosed with epilepsy |
| ● History of frequent or severe headaches |
| ● Diagnosis of tinnitus or hearing problems |
| ● History of fainting, particularly during medical procedures |
| ● Current use of psychoactive medications with potential interactions with TMS |
| ● Excessive alcohol consumption (>10-12 drinks per week) |
| ● Recreational drug use more than once per week |
| ● Diagnosis of color blindness |
| ● Non-Caucasian ethnicity (less than four grandparents of European descent - required for genetic assessments not included in the current data) |

### Experimental design

Data were collected between April 2018 and December 2020 at Monash Biomedical Imaging (MBI), Clayton Campus, Melbourne, Australia. Pre-existing datasets from the GOC (imaging and genetic) and MBBP (cognitive, imaging and genetic) incorporated into the study in accordance with ethical guidelines were collected between February 2014 - August 2017 and September 2018 - December 2020, respectively. An overview of study procedure is listed in Table 3.

**Table 3.** Data acquisition procedure.

| *Days* | *Assessments* | *Details* | *Number of Participants* | *Core Cognitive Domain* |
| --- | --- | --- | --- | --- |
| **Day 0** | Telephone screening<br>(Participant enrolment) | Inclusion | 123 | — |
| **Day 1** | MRI | T1-weighted (MPRAGE) | 123 | — |
|  | Cognitive Test Battery | Bar Task | 120 | Working Memory (Change Detection) |
|  |  | Colour Task | 121 | Working Memory (Change Detection) |
|  |  | Orientation Task | 122 | Working Memory (Continuous Recall) |
|  |  | Symmetry Span | 120 | Working Memory (Complex Span) |
|  |  | Operation Span | 117 | Working Memory (Complex Span) |
|  |  | Backwards Digit Span | 118 | Working memory (Complex Span) |
|  |  | 2-back Task | 117 | Working memory (Updating and Monitoring) |
|  |  | Symbol Search | 120 | Processing Speed |
|  |  | Go/No-Go | 117 | Response Inhibition |
|  |  | Stop-Signal Task | 122 | Response Inhibition |
|  |  | Simon Task | 113 | Response Inhibition |
| **Day 2** | EEG-Rest | Resting state (8 mins) | 121 | — |
|  | EEG-Task | Colour wheel task | 117 | Working Memory (Continuous Recall) |
|  | TMS | Resting motor threshold (RMT) | 123 | — |
|  | TMS–EEG | Prefrontal stimulation | 122 | — |
|  |  | Premotor stimulation | 123 | — |
|  |  | Parietal stimulation | 123 | — |
|  |  | Control stimulation | 120 | — |

New participants completed two assessment sessions, totalling 5-6 hours. Day 1 involved MRI data collection for newly recruited individuals, alongside cognitive and genetic assessments (i.e., saliva sample). Day 2 included EEG data collection both at rest and during task performance, as well as TMS-EEG recordings at rest. Note that data from MRI and genetic assessments are not included in the present dataset.

## SESSION 1

### Magnetic resonance imaging (MRI)

MRI data were acquired on a Siemens 3T Skyra scanner (Siemens Healthineers, Erlangen, Germany). T1-weighted images were acquired using a magnetization-prepared 2 rapid-acquisition gradient-echoes (MP2RAGE) sequence (TR = 5000 ms, TE = 2.98 ms, TI1/TI2 = 700/2500 ms, 1.0 mm isotropic voxel size). MRI was primarily used to guide the position of the TMS coil using stereotaxic neuronavigation.

### Cognitive Assessment

#### Cognitive Test Battery

All cognitive battery tasks were administered electronically by trained research staff and students following standardised procedures. Testing was conducted on a 12.5 inch laptop using Inquisit 5 (Millisecond Software), PsychoPy2 (Peirce, 2007) or Matlab (The MathWorks Inc., 2017b). Each task started with a brief practice version to ensure familiarisation. Newly recruited participants completed the full test battery on day 1, while returning participants undertook an abbreviated version consisting of the remaining tasks which were not completed in previous studies. Tasks were selected to sample cognitive domains commonly distinguished in latent-variable models of working memory and executive function, including working memory storage capacity (change detection/visual array/continuous recall tasks: Bar Task, Colour Task, Orientation Task), working memory storage capacity with distraction (complex span tasks: Symmetry Span, Operation Span, Backwards Digit Span), working memory updating (n-back task), processing speed (Symbol Search Task), and inhibitory control (Go/No-Go Task, Stop-Signal Task, Simon Task) (Engle et al., 1999; Miyake et al., 2000; McAuley and White, 2011).

### Working Memory Storage Capacity

#### Bar Task (BRT)

The Bar Task (fig. 1a) is a change-detection working memory task implemented in MATLAB, adapted from standard visual working memory paradigms (Luck and Vogel, 1997). For this task, participants were presented with sets of oriented (−45°, 0°, 45°, 180°) coloured (red or blue) bars displayed for 100 ms. After a 900 ms delay, a second array appeared, and participants indicated whether any bar had changed orientation or colour by pressing the Z key (“change”) or M key (“no change”) on the keyboard. Each trial was separated by a 200 ms inter-trial interval (ITI). The task comprised 40 trials per condition (4 and 6 loads), yielding 80 total trials (50% change, 50% no change).

**Figure 1.**
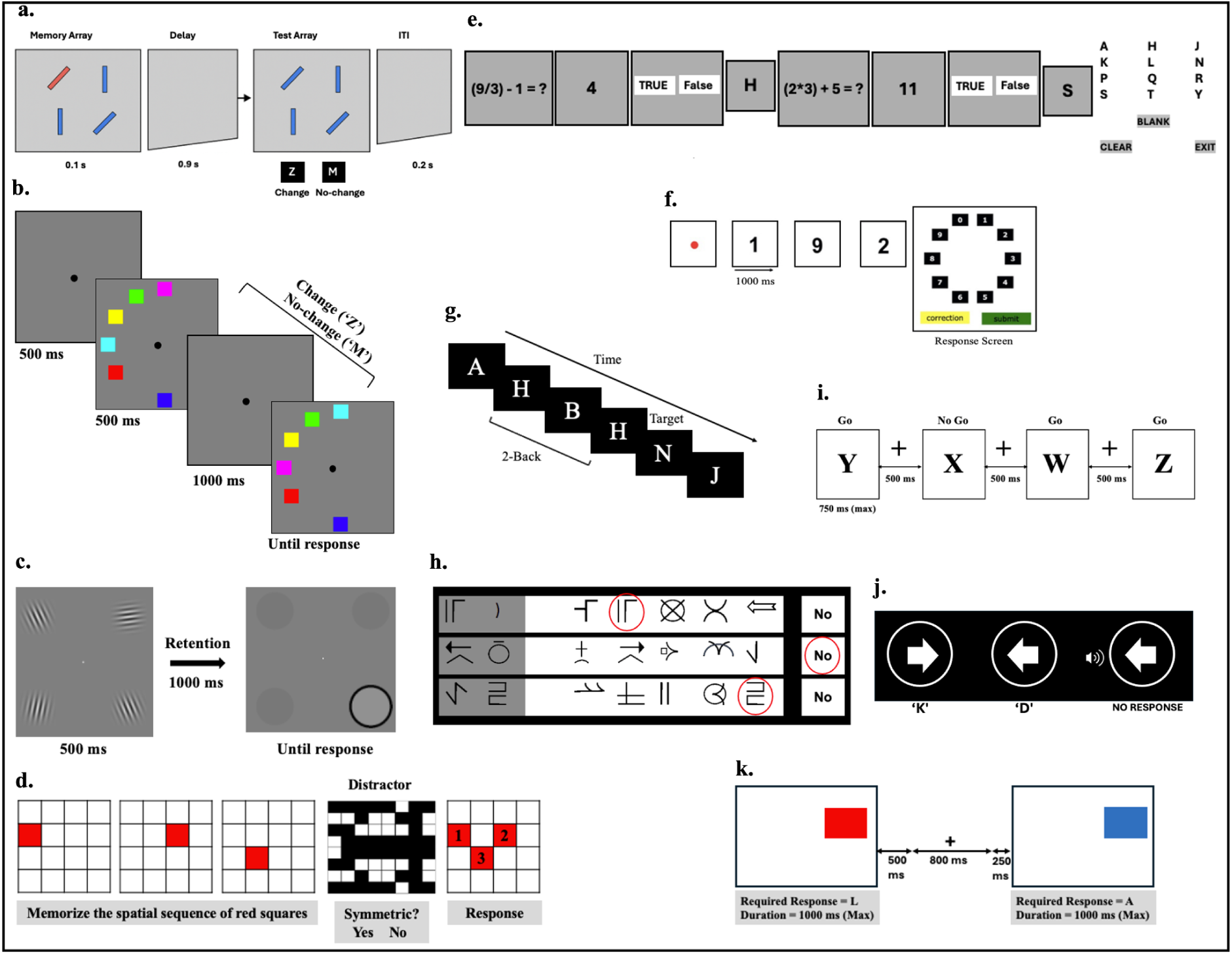
The cognitive battery tasks. (a–k) Representative task structures for each paradigm, illustrating the sequence of events within trials: (a) Bar Task, (b) Colour Task, (c) Orientation Task, (d) Symmetry Span, (e) Operation Span, (f) Backwards Digit Span, (g) n-back task, (h) Symbol Search Task, (i) Go/No-Go Task, (j) Stop-Signal Task, (k) Simon Task.

*Variables recorded:* stimulus and probe features (orientation, colour, load), change condition, participant responses, RT.

#### Colour Task (CLT)

The Colour Task (fig. 1b) (Luck and Vogel, 1997) is a change detection WM paradigm implemented in PsychoPy2. Participants were required to remember the colour and location of a set of coloured squares (500 ms presentation) and indicate whether a subsequent array (1000 ms later) was identical (by pressing the ‘M’ key) or changed (by pressing the ‘Z’ key), with a 500 ms ITI. The task included 13 trials per condition (2-, 4-, and 6-item loads; 42 total trials).

*Variables recorded:* number of items (set size), original stimulus colour, probe stimulus colour, change presence, participant response, correctness, and RT.

#### Orientation Task (ORT)

The Orientation Task (fig. 1c) (Sperling, 1960) was a partial-report WM paradigm implemented in *MATLAB 2017b*. Participants were required to remember the location and orientation of two or four black-and-white circular Gabor patches presented on a grey background (500 ms presentation, 1000 ms retention time) and then reproduce the orientation of a cued target patch using a keypad. The task included 30 trials per condition (2-, 4-item load; 60 total trials).

*Variables recorded:* load condition, target orientation, and orientation error.

### Working Memory Storage Capacity with Distraction

#### Symmetry Span (SYM)

The Symmetry Span task (fig. 1e) (Conway et al., 2005) is a visuospatial complex span measure of working memory capacity and was implemented using *Inquisit 5*. Participants were required to memorize the spatial sequence of red squares presented sequentially within a 4×4 grid while, on alternating trials, making judgments about the symmetry of a pattern of black squares displayed in an 8×8 grid. During each distractor trial, participants indicated whether the black-square pattern was vertically symmetric (“YES”/“NO”) before recalling the red-square sequence in the order of its presentation. The task included 12 trials per condition with two, three or five red squares presented in each.

*Variables recorded:* task phase (show-square, recall-square, show-symmetry, solve-symmetry), load condition (set size), spatial location of squares, recalled square sequence, symmetry stimulus, ground-truth symmetry (symmetrical/asymmetrical), participant response (true/false or grid selection), and RT.

#### Operation Span (OPN)

The Operation Span Task (fig. 1d) (Conway et al., 2005) is a complex paradigm that evaluates WM capacity under dual-task demands and was implemented in *Inquisit 5*. Participants were required to remember a series of alphabetical letters presented for later recall, between alternating trials where they were required to make judgements about the accuracy of answers to mathematical equations. Singular letters (T, L, Q, N, F, H, Y, S, P, K, R, J) appeared in black font on a grey background (1000 ms), which participants were required to later recall in order of appearance by clicking on the letters in the response screen (500 ms interval). During the distractor task, participants were required to evaluate the accuracy of an answer to a mathematical sum by clicking ‘TRUE’ or ‘FALSE’ (5000 ms time out). The task included three trials per load condition (2-7 items) in ascending difficulty (15 total trials).

*Variables recorded:* task phase (show-letter, recall-letter, show-math, solve-math), load condition (set size), letter identity, recalled letter sequence, mathematical problem, ground-truth answer (true/false), participant decision (true/false), correctness, and RT.

#### Backwards Digit Span (BDS)

The Backwards Digit Span task (fig. 1f) (Woods et al., 2011a) was administered in *Inquisit 5* to assess sequence recall. Participants were presented with individual digits (1 s each, black font on white background) and instructed to recall the sequence in reverse order by clicking on-screen digits. Sequence length adaptively increased or decreased depending on participant performance (range = 2–16 digits, 14 total trials).

*Variables recorded:* Backward digit span (maximum sequence length correctly recalled), number of correct trials, number of trials preceding two consecutive errors, and predicted performance score.

### Working Memory Updating

#### 2-Back Task (NBT)

The 2-back working memory task (fig. 1g), based on standard n-back paradigms (Fantz, 1958), was implemented in *PsychoPy2*. Participants were presented a continuous stream of uppercase letters (A–J) presented centrally in white on a black background and instructed to press the down arrow key whenever the current letter matched the one presented two trials earlier. The task contained 130 total trials, including 31 target trials (24%). Each stimulus remained onscreen for 1 s with a fixed ISI of 0.5 s.

*Variables recorded:* probe letter identity, response, correctness, and RT.

### Processing Speed

#### Symbol Search (SBS)

The Symbol Search task (fig. 1k) (Wechsler, 2003) was administered using *Inquisit 5* to measure perceptual speed and visual scanning efficiency. Participants were required to scan rows of characters to determine whether target symbols presented at the start of each row appeared within the search array. Responses were made by clicking on the present character or the “No” button. Stimuli were presented in black font, with target symbols displayed on a grey background and search symbols on a light blue background. Each page contained 10 rows, and participants were given a 2-minute time limit to complete the task.

*Variables recorded:* correctness, total correct sum, and RT.

### Inhibitory Control

#### Go/No-Go (GNG)

The Go/No-Go task (fig. 1i) (Verbruggen and Logan, 2008) assessed inhibitory control and was administered using *Inquisit 5*. Participants were shown letters (“X”, “W”, “Y”, or “Z”) in black on white background and instructed to respond to “W”, “Y”, or “Z” by pressing the spacebar while withholding responses to “X”. Target letters were presented for up to 750 ms, and trials were separated by a 500-ms ISI displaying a fixation cross (+). The task comprised 50 trials per condition (250 total).

*Variables recorded:* correctness, commission errors, omission errors, and RT.

#### Stop Signal Task (SST)

The Stop Signal Task (fig. 1h), a well-established measure of inhibitory control (Logan and Cowan, 1984; Verbruggen et al., 2019), was administered using *Inquisit 5*. Participants were required to respond to left- and right- facing arrows that appeared on the screen one at a time, by quickly pressing the ‘D’ and ‘L’ keys, respectively, except when a tone signalled them to withhold their response. The stop-signal delay (SSD), which was defined as the interval between the disappearance of the target discrimination cue (arrow) and the onset of the stop-signal, was initially set at 250 ms and adjusted by ±50 ms based on performance (range: 50–115 ms). The task comprised three consecutive blocks of 64 trials [192 in total] and lasted approximately 9 min on average.

*Variables recorded:* RT, stop-signal delay, correctness, and successful inhibitions.

#### Simon Task (SMT)

The Simon Task (fig. 1j) (Simon and Wolf, 1963; Bialystok et al., 2004) measured response inhibition and was administered using *Inquisit 5*. In this task, participants were required to quickly respond to red or blue squares presented on the left or right side of the screen by pressing ‘A’ (blue) or ‘L’ (red) key. Prior to the appearance of each square, a fixation cross appeared for 800 ms, followed by a 250 ms blank interval. Then the stimulus square was presented for 1000 ms or until a response was made followed by a 500 ms ITI (100 total trials).

*Variables recorded:* stimulus–response congruency, stimulus colour, stimulus position, response correctness, and RT.

## SESSION 2

### EEG and TMS-EEG session

All participants were seated in a comfortable chair and requested to remain relaxed, with their eyes open and fixated on the screen ahead. Following preparation of the EEG cap, EEG was recorded at rest (8 mins), and then during 4 blocks of the working memory task. The motor hotspot was then determined using TMS and the resting motor threshold determined. Blocks of single pulse TMS were then applied across four conditions (dorsolateral prefrontal cortex, premotor cortex, posterior parietal cortex, shoulder), the order of which was pseudorandomised across participants.

### EEG

EEG activity was recorded with a SynAmps^2^ EEG system (Neuroscan, Compumedics, Australia) from 62 TMS-compatible Ag/AgCl-sintered disk electrodes arranged according to the 10-20 international system in an elastic cap (EASYCAP, Germany). EEG recordings were online-referenced to FPz and grounded to AFz, with electrode placement co-registered to individual structural MRI scans and recorded in real-space using a neuronavigation system (Brainsight^TM2, Rogue Research Inc., Canada). EEG signals were amplified by 1000, low-pass filtered (DC-2000 Hz) and sampled at a rate of 10 kHz. Curry 8 (Neuroscan, Compumedics, Australia) was used to record EEG for offline analysis whilst maintaining skin electrode impedance <5 kΩ.

#### Resting-state EEG

Participants were instructed to remain relaxed, focused ahead with eyes open and to think of nothing in particular while resting-state EEG was recorded (8 mins). The length of resting-state recording was determined to approximately match the time required for each TMS-EEG data collection block (approx ∼8 mins).

#### EEG during WM task

The Continuous Recall Task (fig. 2) (Suchow et al., 2013) is a partial-report WM task administered using Matlab 2017b with concurrent EEG. Participants were presented with a group of coloured squares (2, 4, 6) on a grey background (500 ms, 1000 ms interval) before being prompted to recall the colour of the singular outlined square presented on the next screen by clicking on its position on the colour wheel. There were 100 trials per condition (2, 4, 6) divided into 4 blocks (75 trials per block) (300 total trials).

**Figure 2.**
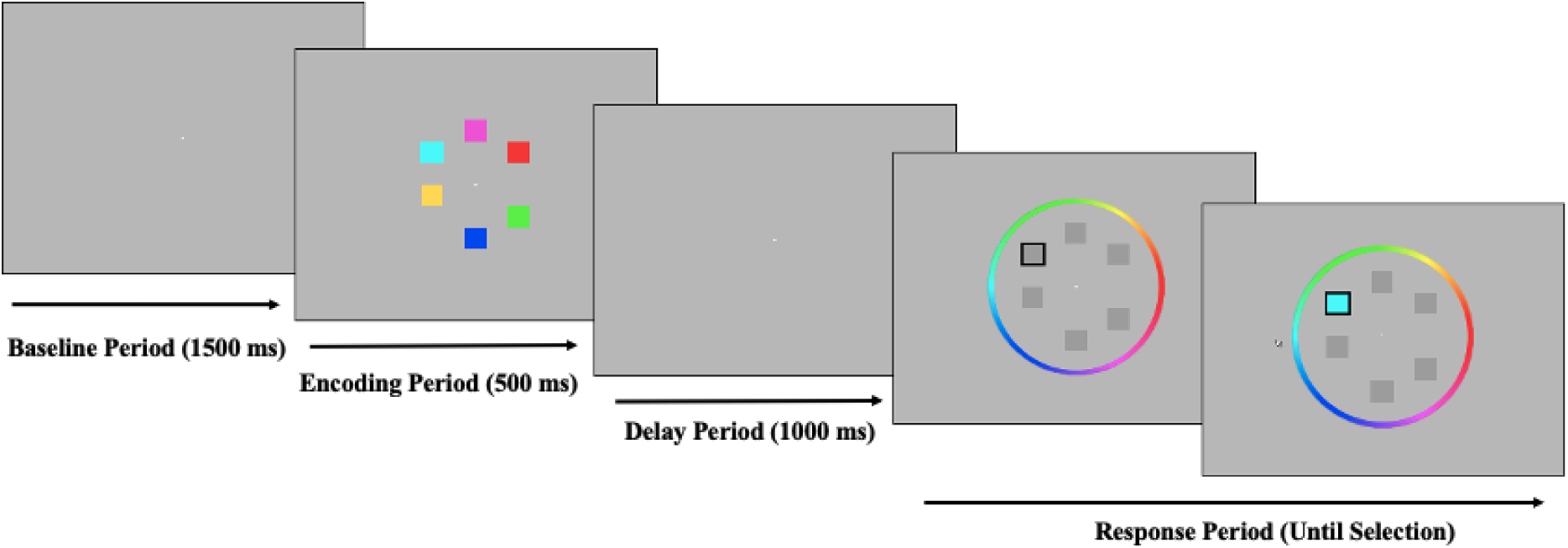
Example display from The Continuous Recall Working Memory Task. Adapted from (Virtue-Griffiths et al., 2025), under the terms of the Creative Commons Attribution 4.0 International License (CC BY 4.0).

*Variables recorded:* Target colour and its degree, non-target colour and its degree, reported colour and its degree, and trial load.

### EMG

Electromyographic (EMG) activity was measured with bipolar surface Ag-AgCl electrodes (4 mm) placed on the right first dorsal interosseous (FDI) muscle in a belly-tendon montage (2 cm distance), grounded at the right-hand dorsum. EMG signals were amplified by 1000, band-pass filtered (10-1000 Hz), sampled at a rate of 5 kHz, epoched around the TMS pulse (-300 to 500 ms) and displayed online on the computer screen.

### TMS

TMS was administered using a Magventure CB60 figure-of-eight coil attached to a MagPro X100 with Option stimulator (MagVenture, Denmark). The stimulator was controlled via the MAGIC toolbox (Habibollahi Saatlou et al., 2018) to deliver biphasic pulses to the cortex in an anterior-posterior and posterior-anterior direction. A stereotactic neuronavigation system was used to guide coil placement on the basis of individual neuroanatomy (Brainsight, Rogue Research, Montreal, QC, Canada). First, resting motor threshold (RMT) was determined by: 1) finding the motor ‘hotspot’ defined as the coil position over left primary motor cortex resulting in the largest motor evoked potentials in right FDI muscle, and 2) finding the minimum intensity to elicit a motor evoked potential >0.05 mV in at least 5/10 trials while wearing the EEG cap. Next, four blocks of stimuli were delivered targeting four different locations: 3 cortical and 1 control (shoulder) location. In total, 120 pulses were delivered with 4 - 6s jittered inter-pulse intervals per block. Targeted cortical regions were within the left hemisphere and included the dorsolateral prefrontal cortex (x = -46, y = 33, z = 34; Brodmann area 9), premotor cortex (x = -23, y = 4, z = 63; Brodmann area 6) and posterior parietal cortex (x = -26, y = -69, z = 63; Brodmann area 7). Stimulation sites were chosen to overlay peaks in haemodynamic activity identified following a meta-analysis of 1091 fMRI studies assessing working memory generated by Neurosynth (Yarkoni et al., 2011). The target coordinates were adjusted for each individual to ensure positioning of the coil over the crown of the nearest gyrus. Coil placement for cortical stimulation was tangential to the left side of the head, with the coil handle positioned at an approximate 90-degree angle to the underlying gyrus. Small additional adjustments to coil angle were made to minimise sensation of the pulse on the scalp via feedback from the participants. Stimulation intensity was determined by calculating the RMT adjusted for differences in coil-to-cortex distance (RMT_adj_) for each site (Stokes et al., 2005, 2007). RMT_adj_ was calculated at each location by: 1) calculating the distance between the centre of the coil and the target location on the cortex (in mm); 2) multiplying the distance by a scaling factor of 2.8; and 3) adding the value to the RMT. The first 94 participants received TMS at 100% RMT_adj_, while the final 29 received TMS at 120% RMT_adj_. The initial stimulation intensity of 100% RMT_adj_ was chosen for consistency with studies targeting the motor cortex which used this intensity to minimise sensory input associated with stimulation. However, stimulation intensity was adjusted for the last 29 participants following preliminary analyses which indicated a higher signal-to-noise ratio for TMS-evoked potentials when using higher TMS intensities. A control condition was also incorporated to mimic auditory and somatosensory effects elicited by TMS, where 120 pulses were delivered to the left acromioclavicular joint over the shoulder (Herring et al., 2015; Biabani et al., 2019). Coil positioning and intensity for shoulder stimulation was modified relative to participants’ reported comparisons of local sensations produced by the TMS coil at the strongest cortical site. Coil angle and orientation was further modified according to visual inspection and participant feedback to reduce pulse transmission down the arm and hand. Stimulation order between blocks was pseudo randomised for each participant.

Additional control measures were implemented during stimulation to further minimise multisensory effects of TMS which were considered effective at the time the experiments commenced (Ilmoniemi et al., 2015). A thin sheet of foam (∼3 mm) was attached to the underside of the coil to reduce sensory input from coil vibration and bone conduction, while white noise was played through earphones to minimise auditory activation from the acoustics of the TMS click. White noise volume was increased to a level where stimulation was no longer perceivable to participants or their upper threshold of comfort for acoustic stimuli was reached. Following each block of stimulation, participants completed four questions on a 10-point Likert scale rating the sensory experiences of stimulation including:

1. How uncomfortable were the TMS pulses?
2. If painful, how intense was the pain from the TMS pulses?
3. If there were any twitches, how strong were the muscle twitches from the TMS pulse?
4. How well did the white noise cover the sound of the TMS pulse?

Despite the sensory masking process, the majority of participants reported that they were still able to distinguish the clicking sound and scalp sensation while receiving stimulation, particularly for stimulation sites in the frontal cortex (Biabani et al., 2024).

### EEG pre-processing

Prior to public release, EEG data were partially pre-processed in MATLAB (R2023a, The MathWorks, USA) using EEGLAB (Delorme and Makeig, 2004) and TESA (Rogasch et al., 2017) to reduce file size. All working-memory EEG blocks were concatenated into a single dataset for each participant. For continuous TMS-EEG recordings, the TMS pulse artefact was removed (−1 to 5 ms) and then reconstructed via cubic interpolation. All EEG datasets were then down-sampled to 1000 Hz. MRI data is not publicly available as part of this dataset. This is due to reasons including public release not being stated as the intent of MRI data use at the time of collection for prior studies, as well as concerns of confidentiality in the ability to reverse defaced NIFTI images (Schwarz et al., 2021; Clunie et al., 2025; Gao et al., 2025).

### Data Record

The data can be accessed from the OpenNeuro public repository (access number: ds008037). This OpenNeuro release is organised according to the Brain Imaging Data Structure standard (Gorgolewski et al., 2016), including the EEG-BIDS extension (Pernet et al., 2019), and the emerging NIBS-BIDS/BEP037 recommendations (Bertazzoli et al., 2026), where applicable.

### Technical Validation

Datasets were manually checked for missing or corrupt data prior to their inclusion into the database. Subsequent quality checks were done on the EEG and behavioural data as follows:

#### Resting-state EEG data quality

The quality of resting-state EEG data was evaluated by examining the spectral characteristics of the recordings. We first preprocessed the data using an automated cleaning pipeline (Reduction of Electroencephalographic Artifacts (RELAX) (Bailey et al., 2022, 2023)) implemented within EEGLAB (Delorme and Makeig, 2004). Briefly, data were bandstop (47–53 Hz) and bandpass (0.25–80 Hz) filtered using zero-phase Butterworth filters. Noisy channels were then identified and removed using the PREP pipeline (Bigdely-Shamlo et al., 2015), followed by two rounds of multi-channel Wiener filtering (MWF) to reduce eye-movement, muscle, and drift-related artefacts. Residual artefacts were identified using the ICLabel independent component analysis (ICA) classifier (Pion-Tonachini et al., 2019) and the artifactual components were subsequently attenuated using wavelet-enhanced ICA (wICA) implemented with the FastICA symmetric algorithm. Rejected electrodes were interpolated using spherical interpolation and the data were re-referenced to the common average. Final cleaned datasets were retained as continuous EEG data for subsequent analyses. The power spectral density (PSD) was estimated for each individual and each channel using Welch’s method. Group-level spectra showed an expected 1/f like decay across frequencies with a clear peak in alpha range (around 10 Hz; fig. 3A). The alpha power was maximal around occipitoparietal regions, consistent with the well-established neurophysiological pattern observed at rest (Jensen and Mazaheri, 2010); fig. 3B). We also examined the integrity of channel-level data across individuals and frequencies and observed broadly homogeneous power distributions across participants (fig. 3C). There was no evidence of systematically noisy or flat channels, or globally abnormal recordings, indicating stable signal quality across the dataset.

**Figure 3.**
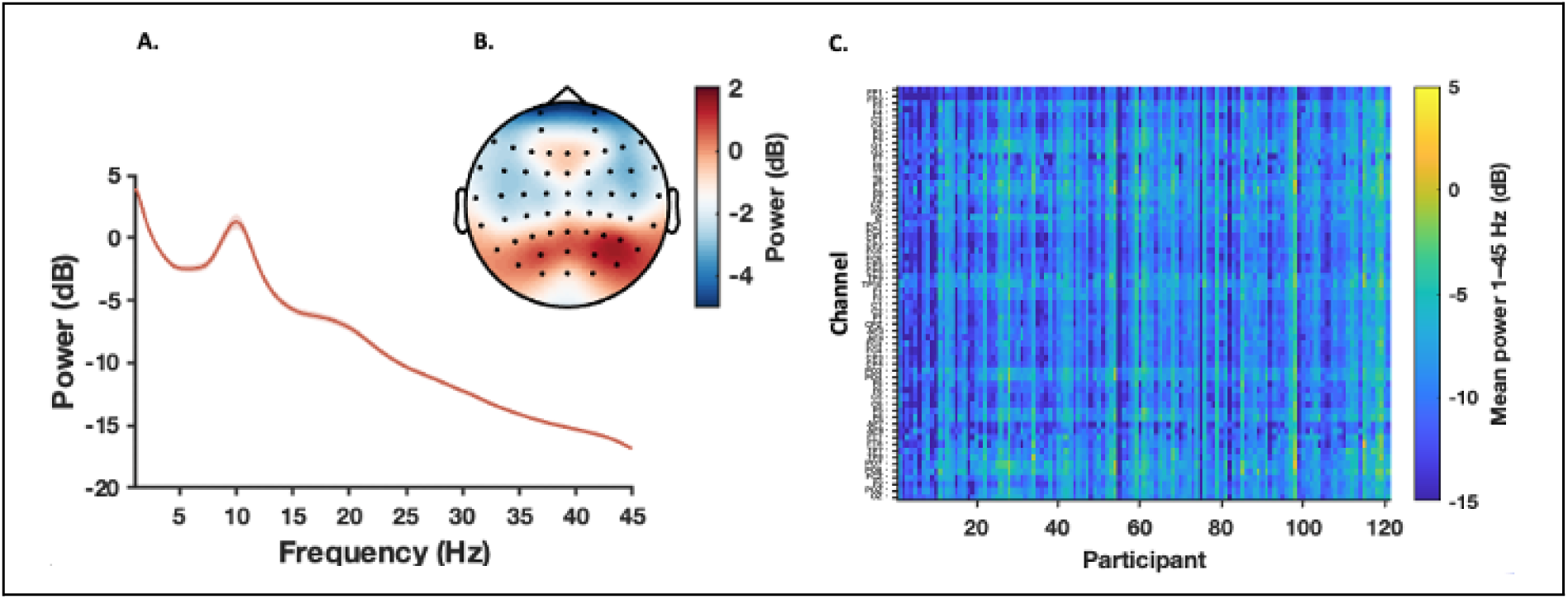
Quality assessment of resting-state EEG data. (A) Mean spectral power of EEG signals recorded at occipito-parietal regions (O1, Oz, O2, POz, Pz), showing the 1/f-like decay with a pronounced peak at alpha band (8-12 Hz). (B) Spatial distribution of alpha power centred around occipito-parietal regions. (C) Channel-by-participant heatmap of broadband power spectrum (1–45 Hz), indicating consistent signal quality across participants and channels, with only minor and localized deviations, and no evidence of systematic corrupted recordings.

#### WM data quality (EEG and performance)

The quality of task-EEG data was assessed via visualisation of time-locked responses and channel-level signal characteristic. The raw recordings underwent the same cleaning procedure as the resting state data. Subsequently, grand-average event-related potentials (ERPs) were computed separately for each working memory load condition (2, 4, 6) and visualised at selected regions known to contribute to the neural dynamics of VWM (Prefrontal-F3; Premotor-FC1; Parietal-P3; (Miller and Cohen, 2001; Curtis and D’Esposito, 2003; Todd and Marois, 2004; Harrison and Tong, 2009). We found canonical time-locked components in the signals which were showing systematic modulation by task load indicative of physiologically meaningful neural activity (fig. 4A). To examine the integrity of the recorded signals across channels and participants, we visualised the root-mean-square (RMS) of the amplitude using a heatmap (fig. 4B). Overall, the recorded potentials were consistent across the dataset, with no indication of systematically flat or excessively noisy channels or any global abnormality.

**Figure 4.**
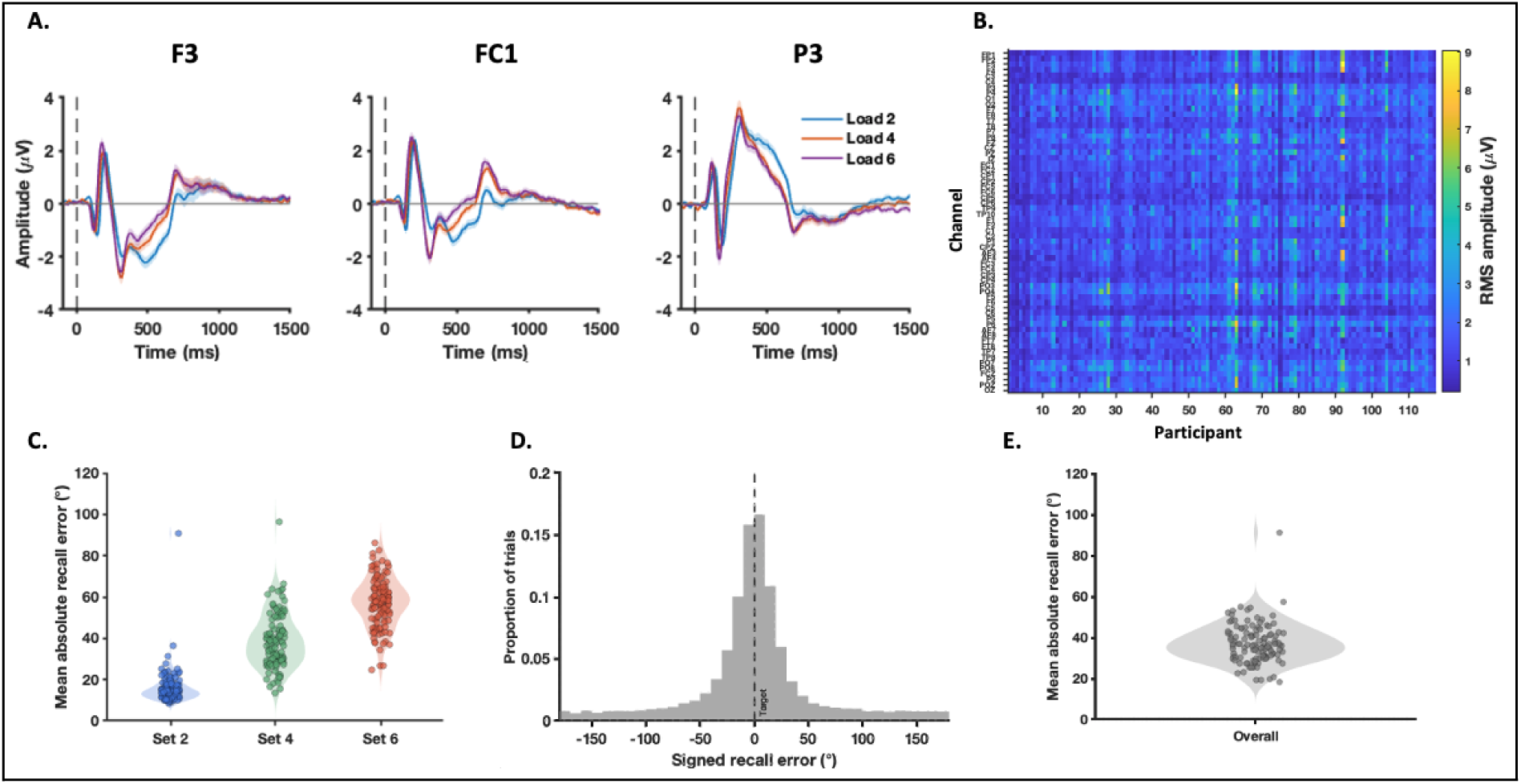
Quality assessment of the Visual working memory EEG and task performance data. (A) Grand average ERPs (mean ± SEM) recorded at Prefrontal (F3), Premotor (FC1), and Parietal (P3) regions at the three task loads. All ERPs exhibit clear canonical components with region-specific and load-sensitive behaviour. (B) Channel-by-participant heatmap of RMS amplitude showing little evidence for excessive noise. (C) Distribution of mean absolute recall error across participants at the three task loads confirming sensitivity of performance to task difficulty. (D) Histogram plots showing the distribution of signed recall errors (-180° to 180°) across trials. The clustered values around 0° (indicative of the target colour being chosen) argues against randomness of the responses. (E) The average of absolute recall error across loads showing normal distribution, which supports a biological behavior inside the population. Circles inside violin plots represent values for each individual.

For the quality control of performance data we examined the distribution of recall errors (difference between reported and target colours) and their sensitivity to the difficulty of the task (set size) (Zhang and Luck, 2008; Suchow et al., 2013). The recall errors increased with set size, consistent with the expected load-dependent constraints on working memory precision (fig. 4D). The signed recall errors (-180° to 180°) showed bell-shape distribution peaking around the target colour indicating unbiased and systematic response (fig. 4D). The overall absolute errors (0 to 180°) were also well below the ∼90° expected under random responding and tightly clustered across individuals (fig. 4E). Although one participant showed elevated condition-specific errors exceeding the benchmark (sub-012; set-size 2, 90.83°; set-size 4, 96.53°; Fig. 4C). The present assessments confirm the stability and interpretability of both EEG and performance datasets for further analyses.

#### TMS-EEG data quality

The interpretability and suitability of the TMS–EEG data have been established in our recent work (Biabani et al., 2024). TMS–EEG data were preprocessed using the validated pipeline described in (Rogasch et al., 2014, 2017) and detailed in (Biabani et al., 2019). Briefly, data from all conditions were concatenated to ensure identical preprocessing across conditions and underwent epoching around the TMS pulse (−1000 to 1000 ms), baseline correction (−500 to −10 ms), removal of the TMS pulse artefact (−2 to 15 ms) followed by cubic interpolation, and downsampling to 1000 Hz. Data were then visually inspected for artefacts, and TMS-induced muscle and decay artefacts were removed using FastICA. This was followed by band-pass (1–100 Hz) and band-stop (48–52 Hz) filtering using a zero-phase Butterworth filter (order = 4), and a second ICA to remove residual artefacts (e.g., eye blinks, channel noise). Finally, rejected channels were spherically interpolated, and the data were re-referenced to the common average. Figure 5 (adapted from (Biabani et al., 2024)) illustrates the characteristics of the processed TEPs to both high (120% RMT_adj_) and low (100% RMT_adj_) stimulation intensities. As shown, TMS at both intensities produced clear canonical peaks in EEG recordings whose magnitude varied as a function of stimulation condition. Consistent with the TMS-EEG literature, early responses (<60 ms) showed site-specific spatiotemporal variation, whereas later peaks (N100/P200), exhibited a broader frontocentral topography, consistent with sensory evoked potentials (Freedberg et al., 2020; Rogasch et al., 2020; Ahn and Fröhlich, 2021).

**Figure 5.**
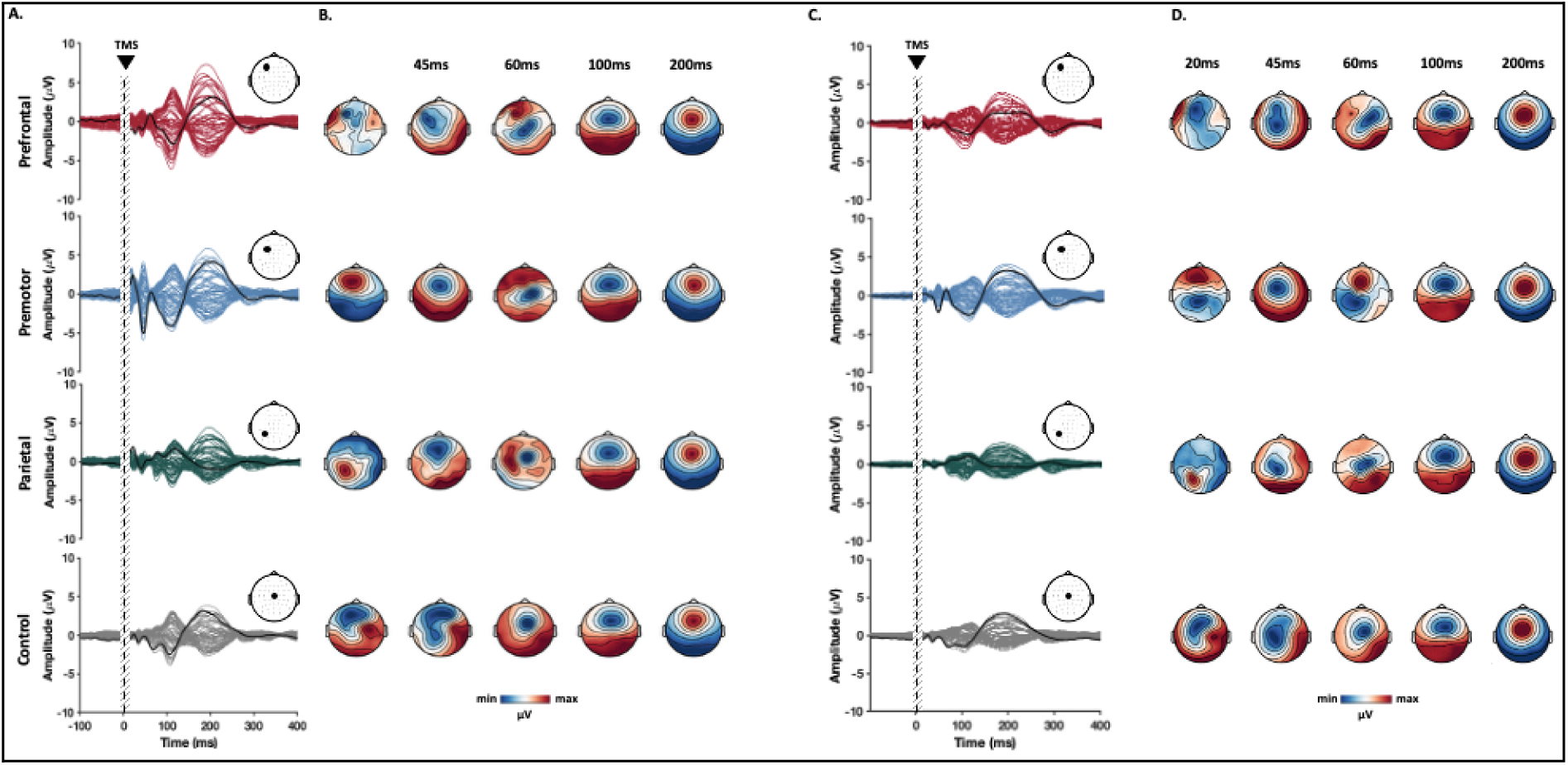
Spatiotemporal distribution of TMS-evoked potentials showing canonical components and condition sensitivity. (adapted from (Biabani et al., 2024), under the terms of the Creative Commons Attribution 4.0 International License (CC BY 4.0)). (A-B) TEPs following high intensity stimulation (120% RMT_adj_; n= 29). (C-D) TEPs following low intensity stimulation (100% RMT_adj_; n= 94). (A, C) Temporal pattern of group-average TEPs recorded at each electrode. The highlighted trace in the butterfly plot and the corresponding electrode on the scalp map denote representative electrodes for each condition (Prefrontal: F3; Premotor: FC1; Parietal: P3; Control: CZ). Grey vertical bars indicate interpolated time windows excluded from analysis. (B–D) Topographical maps show the scalp distribution of potentials at representative TEP peak latencies.

### Cognitive Test Battery Data

For the quality check of the cognitive battery data we examined response distributions and task-specific performance signatures to ensure data consistency and interpretability.

The change detection tasks were evaluated to determine whether they meaningfully challenged working memory storage capacity. For the Bar and Colour tasks we quantified sensitivity by calculating *d’* value, which is the ability of an individual to distinguish between a signal (stimulus) and noise (no stimulus) (Stanislaw and Todorov, 1999). In both tasks the majority of participants showed d’ > 0 across loads confirming the ability to distinguish signals from noise (fig. 6A,C). Consistently, the level of accuracy was reliably above chance (50%) for most participants in both tasks and decreased with task difficulty (fig. 6B,D). The quality of orientation task data were checked based on the distribution of recall errors. As depicted in Figure 6F, the median absolute error values were low, tightly distributed across individuals at low values (<30°) and consistently increased with task difficulty. Overall, the quality check of the detection tasks data, confirm consistent recalls across participants and good task engagement with little evidence of random responding or performance heterogeneity.

**Figure 6.**
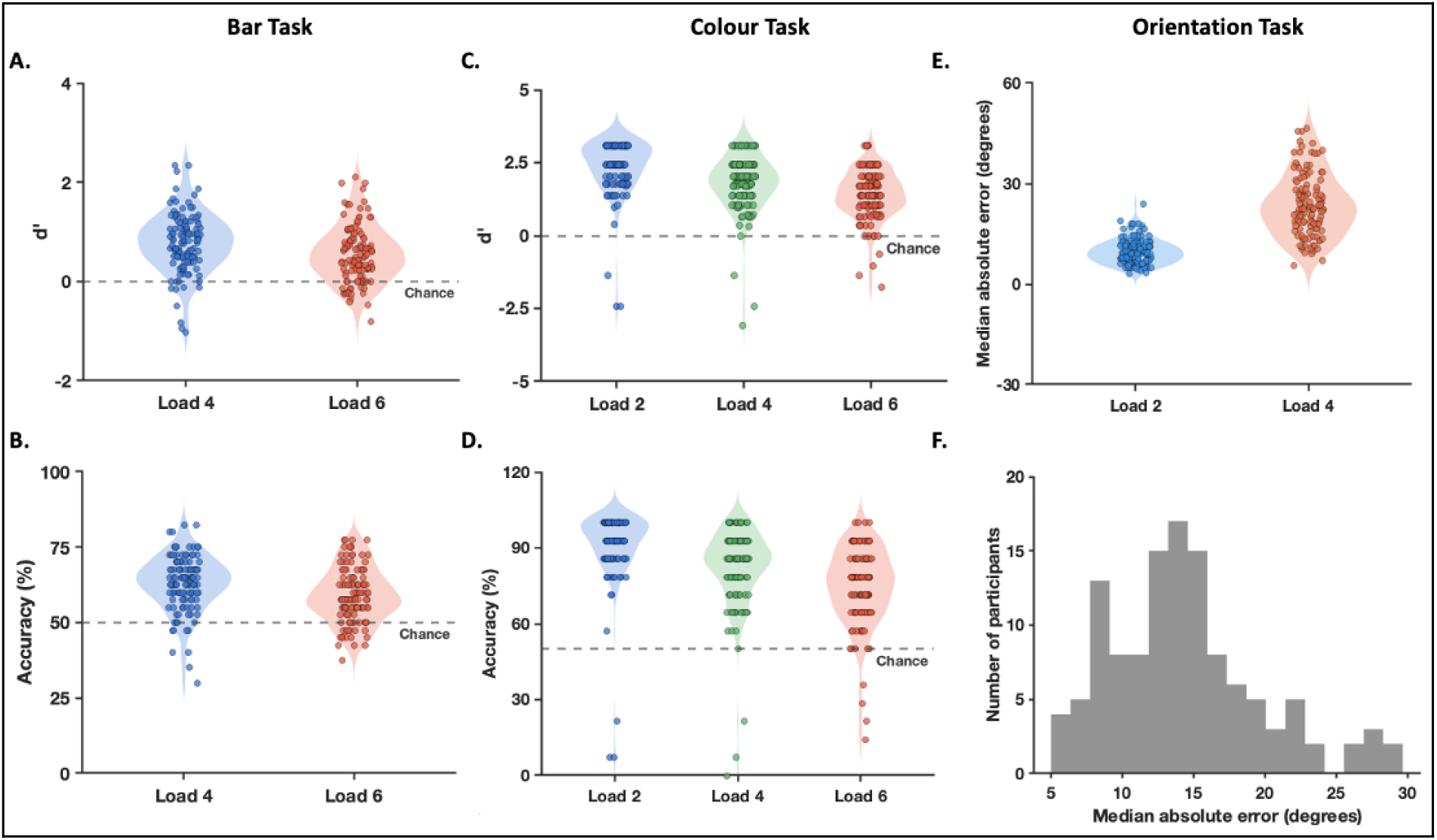
Quality assessment of the Change Detection task performance data. (A-B) Bar task performance. Sensitivity (d′; A) and accuracy (B) are shown separately for memory loads 4 and 6. Performance was reliably above chance (d′ = 0; accuracy = 50%, dashed line), without ceiling effects, and decreased with increasing load. (C-D) Colour change detection task. Sensitivity (d′; C) and accuracy (D) are shown for loads 2, 4, and 6. Performance remained above chance across all loads and decreased systematically with increasing set size. (E–F) Orientation reproduction task. Median absolute error increased with load (E). Histogram of participant-level median absolute error (F) showing unimodal distribution centred at low values. Circles inside violin plots represent values for each individual.

For the Symmetry Span and Operation Span tasks, we evaluated the data based on both memory and processing components of the paradigms. For the Symmetry Span task we first quantified the memory performance using a partial-credit item score, defined as the proportion of square locations recalled in the correct serial position for each memory set (Conway et al., 2005). We observed lower accuracy and greater variability at larger set sizes (Fig. 7A), indicating that the task manipulation increased visuospatial working-memory demands.

**Figure 7.**
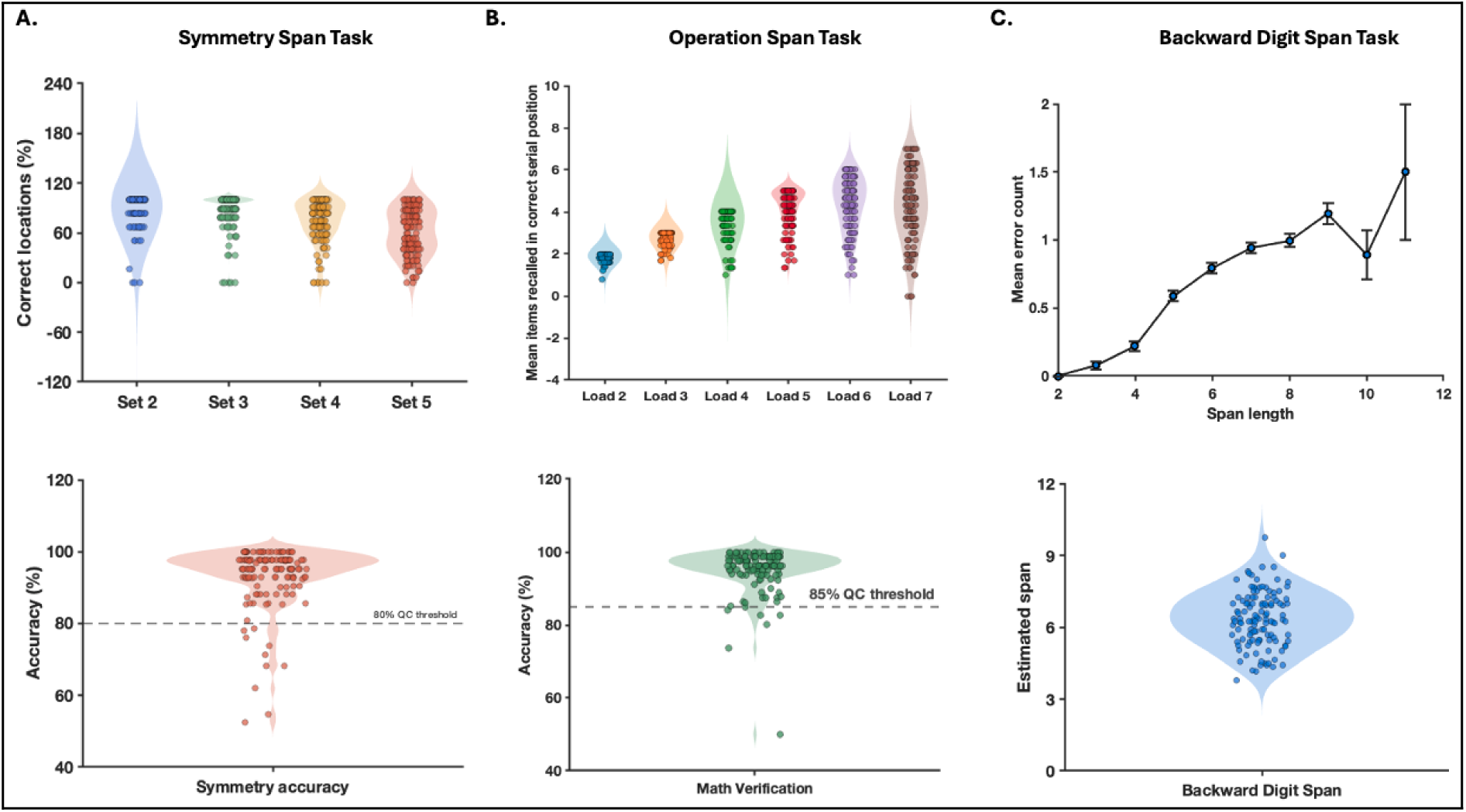
Quality assessment of complex span task performance. (A) Symmetry Span task: the upper plot illustrates partial-credit spatial recall performance by set size, and the lower plot represents the distribution of accuracy on the concurrent symmetry-judgement task. (B) Operation Span task: the upper violin plots depict the mean number of items recalled in the correct serial position across loads, and the lower plot shows the distribution of accuracy on the concurrent mathematical verification task. Dashed lines (A-B; lower plots) indicate the 85% accuracy benchmark for the concurrent processing components of the span tasks. (C) Backward Digit Span task: the upper line graph plots the mean±SEM recall error count across span lengths, and the lower violin plot shows the distribution of estimated span values. Violin plots show participant-level distributions with each dot representing an individual participant.

Furthermore, accuracy on the symmetry-judgement task generally exceeded the quality-control threshold (85% accuracy (Conway et al., 2005)) confirming the engagement of the participants in the concurrent processing task (fig. 7B). Together, these results demonstrate the expected behavioural profile of a complex span task: load-dependent reductions in spatial recall alongside reliable concurrent-task engagement.

Operation span task performance was assessed using both partial-credit (Conway et al., 2005) and absolute recall measures, together with accuracy on the concurrent mathematical verification task (fig. 7B). Partial-credit recall by load showed a systematic effect of memory load, with near-ceiling recall at lower loads (2-3) and a gradual decline in partial-credit performance as load increased (5-7). As expected (Conway et al., 2005), individual variability increased with load and the scores moved towards a normal distribution. This confirms that the task successfully challenged the executive attention and cognitive control of a wide range of working memory capacity. These quality assessments were supplemented by symmetry judgement accuracy, which exceeded the commonly-used 85% benchmark, confirming reliable engagement with the processing component of the task.

For the Backward Digit Span task, we examined the load-dependency of the performance as well as the distribution of the participant-level span estimates (fig. 7C). As shown, the error counts increased with sequence length indicating that the task became progressively more challenging as working-memory demands increased. Performance variability increased with sequence length likely reflecting the smaller number of trials/participants reaching those levels of difficulty. The estimated span values show coherent distribution within the expected range for healthy adults (Woods et al., 2011b), with no evidence for marked floor or ceiling effects. Overall, these findings confirm the interpretability of the present dataset for measuring working-memory capacity.

For working memory updating (n-back) and processing speed (Symbol Search) tasks, we first looked at the distribution of median RTs to identify any implausible or extreme response patterns. For both tasks, RT distributions were unimodal and fell within plausible ranges for the respective paradigms (n-back task : ∼0.45-0.65 s (Hautus et al., 2021); Symbol Search task: ∼2-4 s (Wechsler, 1955; Salthouse, 1996)), with no evidence for anticipatory or random responding (i.e., excessively fast responses)(fig. 8A,C). To evaluate task engagement, we evaluated d′ for the n-back task and found the values above chance (0) confirming reliable discrimination between target and non-targets (fig. 8B). For the Symbol Search task, we examined the distribution of total correct responses and found a unimodal distribution with no evidence of ceiling or floor effects (no clustering near maximum or minimum values) (fig. 8D).

**Figure 8.**
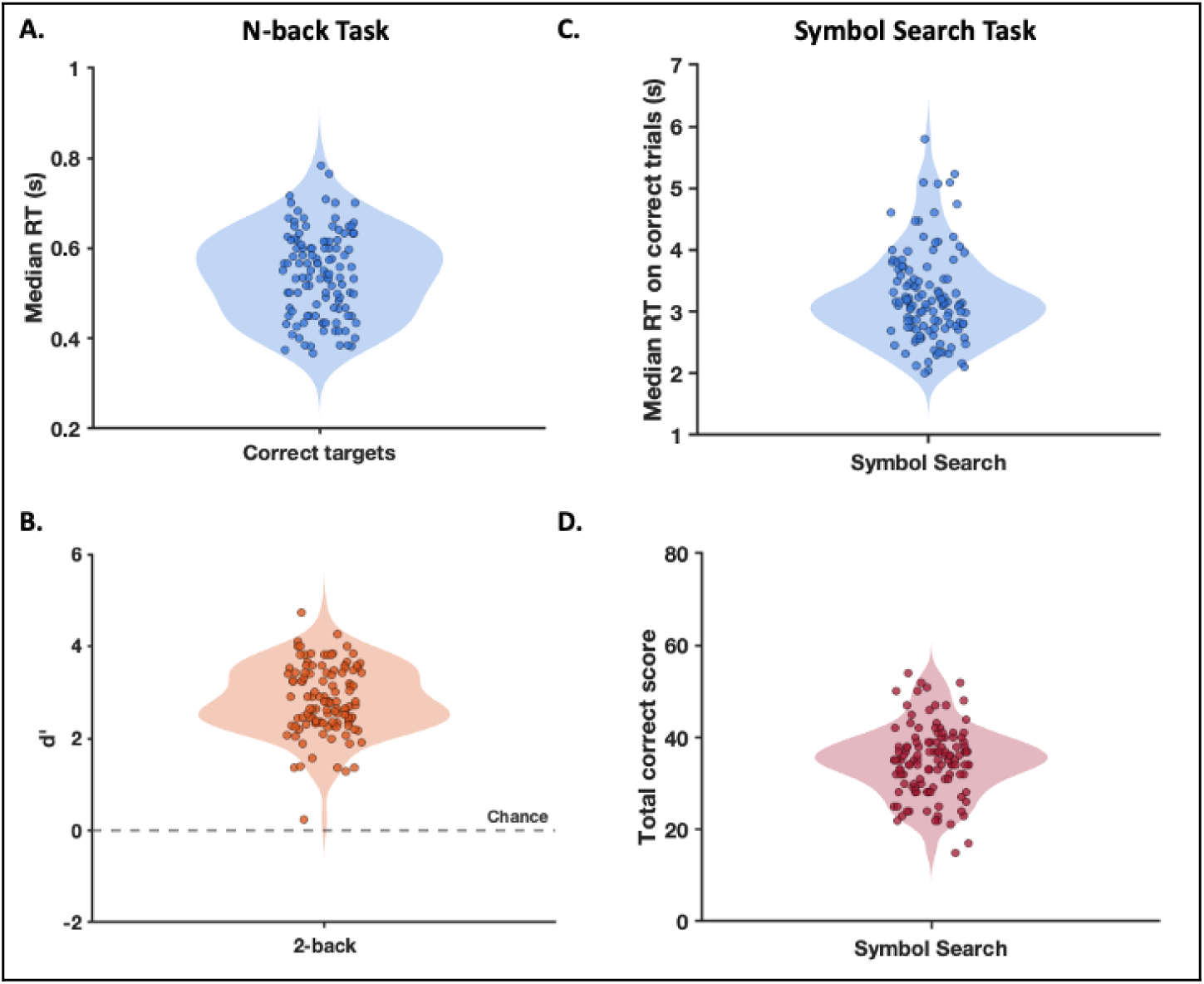
Quality assessment of the N-back and Symbol Search task performance data. (A-B) The distribution of response time (Median RT; A) and sensitivity (d′; B) for the N-back task. (C-D) The distribution of response time from correct trials (Median RT; C) and accuracy (Total correct score; D) for the Symbol Search task. RTs were unimodally distributed and fell within a plausible range for the respective paradigms. Sensitivity values (*d’*) were consistently above chance (d′ = 0; dashed line). Total correct scores in the Symbol Search task showed a unimodal distribution without floor or ceiling effects.

To assess the data from the Go/No-Go task we compared the accuracy of responses between Go and No-Go trials (fig. 9A). Consistent with the expected inhibitory demands, we found less accuracy in No-Go trials, where participants must withhold a prepotent response (Verbruggen and Logan, 2008; Wright et al., 2014). Moreover, response time values on correct Go trials fell within the several-hundred millisecond range commonly reported in healthy young adult Go/No-Go paradigms (Wright et al., 2014). For the Stop-Signal task, we first evaluated whether the behavioural data follow the assumptions of the independent race model (Verbruggen et al., 2019). As expected, responses on failed-stop trials were faster than those on Go trials, indicating that failed inhibitions primarily occurred when the go process finished before the stop process. We also evaluated whether the adaptive staircase procedure effectively calibrated the difficulty of the task across participants (fig. 9B). Importantly, Stop success rates centred around 50% confirming that the task was well-calibrated to generate valid inhibition data (Verbruggen et al., 2013). In the Simon task (fig. 9C), participants demonstrated consistently high accuracy across both congruent and incongruent trials (with higher accuracy for the congruent trials, as expected), which confirms their engagement with the task. To assess the presence of Simon interference effect, we compared the response times between congruent and incongruent trials. As expected, responses were slower on incongruent trials indicating a positive Simon effect across participants (Simon and Rudell, 1967). Together, the observed patterns across the inhibitory-control tasks data indicate that these tasks elicited the expected demands on response inhibition, as well as, response-conflict processing supporting the suitability of the data for subsequent analyses.

**Figure 9.**
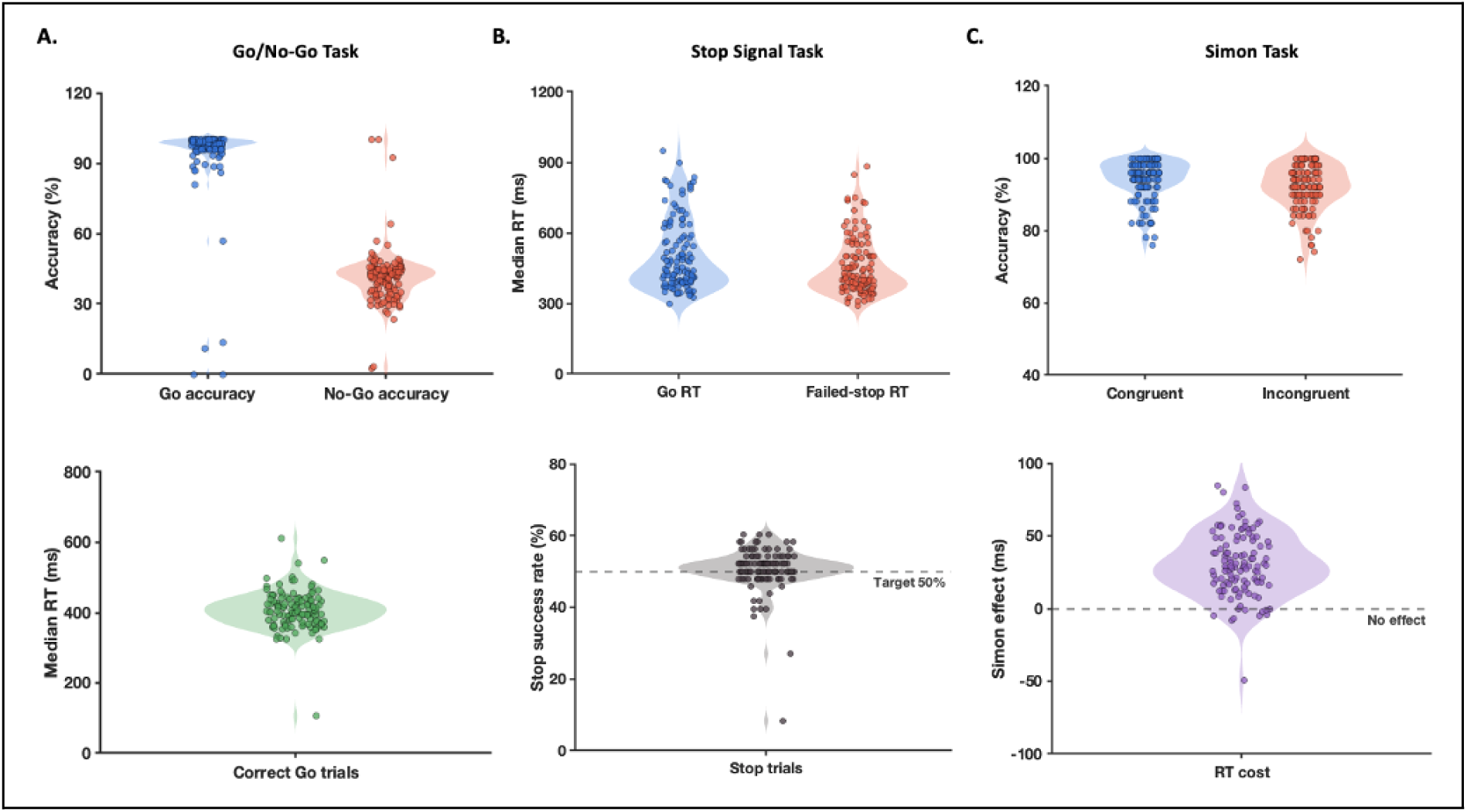
Quality assessment of inhibitory control task performance. (A) Go/No-Go task: the upper plot compares response accuracy between Go and No-Go trials, and the lower plot shows the distribution of response times on correct Go trials. (B) Stop-Signal task: the upper plot compares response times for Go trials and failed-stop trials, and the lower plot shows stop success rates. The horizontal dashed line indicates the 50% target expected from the adaptive staircase procedure. (C) Simon task: the upper plot compares accuracy between congruent and incongruent trials. The lower plot shows the Simon effect (positive; slower response in incongruent trials). The dashed line indicates no Simon effect. Violin plots show participant-level distributions with each dot representing an individual participant.

## Data Availability

The dataset described in this Data Descriptor is available from OpenNeuro under accession number ds008037 [doi:10.18112/openneuro.ds008037.v1.0.0]. The repository contains the de-identified behavioural, EEG, and TMS–EEG data organised according to the Brain Imaging Data Structure (BIDS), together with the associated metadata files required to interpret the dataset.

## Code Availability

Open-source code for the data-processing pipelines and quality-control procedures associated with this dataset is available at : https://github.com/ManaBiabani/GWM_dataset_processing_QC.

## Acknowledgements

This research was supported by the Australian Research Council (ARC) DP170100738 (NCR, AF, ZH, MY), DE180100741 (NCR) and FT210100694 (NCR).

